# Distinct Neural Representations of Hunger and Thirst in Neonatal Mice before the Emergence of Food- and Water-seeking Behaviors

**DOI:** 10.1101/2024.09.22.614378

**Authors:** David C. Wang, Yunming Wu, Conor Mehaffy, Leslie A. Espinoza-Campomanes, Fernando Santos-Valencia, Kevin M. Franks, Liqun Luo

## Abstract

Hunger and thirst are two fundamental drives for maintaining homeostasis and elicit distinct food- and water-seeking behaviors essential for survival. For neonatal mammals, however, both hunger and thirst are sated by consuming milk from their mother. While distinct neural circuits underlying hunger and thirst drives in the adult brain have been characterized, it is unclear when these distinctions emerge in neonates and what processes may affect their development. Here we show that hypothalamic hunger and thirst regions already exhibit specific responses to starvation and dehydration well before a neonatal mouse can seek food and water separately. At this early age, hunger neurons drive feeding behaviors more than do thirst neurons. *In vivo* Neuropixels recordings in dehydrated and starved neonatal mice revealed that maternal presentation leads to a relative increase in activity which is suppressed by feeding on short timescales, particularly in hypothalamic and thalamic neurons. Changes in activity become more heterogeneous on longer timescales. Lastly, within neonatal regions that respond to both hunger and thirst, subpopulations of neurons respond distinctly to one or the other need. Combining food and water into a liquid diet throughout the animal’s life does not alter the distinct representations of hunger and thirst in the adult brain. Thus, neural representations of hunger and thirst in mice become distinct before food- and water-seeking behaviors mature and are robust to environmental changes in food and water sources.

## INTRODUCTION

Hunger and thirst elicit feeding and drinking to maintaining energy and water homeostasis, respectively. Neural circuits regulating hunger and thirst are well distinguished in the adult mammals. For hunger, a well-studied hypothalamic region is the arcuate nucleus (ARC)^1^. Agouti-related protein (AgRP)-expressing neurons in ARC are activated by starvation (we refer to these as hunger neurons hereafter) and necessary for feeding in adult mice^2,3^. Stimulating AgRP hunger neurons is sufficient to drive feeding in sated animals^4^. AgRP neurons project to a wide array of regions both within and outside of the hypothalamus, which have been described in regulating various aspects of feeding behavior^5–8^. For thirst, well-studied brain regions include the subfornical organ (SFO), vascular organ of the lamina terminalis (OVLT), and median preoptic nucleus (MnPO)^1^. In thirsty animals, specific excitatory neuron types in the SFO and OVLT that express an angiotensin receptor (Agtr1a) are activated by an increase in blood osmolarity and in the level of hormone angiotensin II^9,10^, and project to and activate specific population of Agtr1a+ MnPO excitatory neurons (for simplicity, we refer to these dehydration-activated SFO, OVLT, and MnPO neurons as thirst neurons hereafter). Stimulating SFO or MnPO thirst neurons is sufficient to drive drinking behavior in hydrated animals^9–12^.

While hunger and thirst circuits have been well characterized in the mature brain, their functions in neonatal stages are much less clear. Neonatal mammals do not have sources of food and water outside of maternal milk, and thus obtain both nutrition and hydration through the same source. Given that mice do not independently seek food or water at this early developmental stage, it is unclear if neonatal thirst- or hunger-regulating hypothalamic circuits respond distinctly to these needs or if they overlap in their response given their common behavioral output is increased attachment to the mother and suckling for milk. Furthermore, some studies have suggested that instead of promoting need-specific behaviors, neonatal hypothalamic populations such as the AgRP hunger neurons drive maternal-seeking behaviors rather than milk consumption^13,14^. Other studies have suggested that hunger-regulating hormones primarily play a role in the maturation of hypothalamic circuits in neonatal stages rather than in the direct regulation of feeding^15–18^. Thus, it is unclear whether adult hunger and thirst neurons are distinct in their responses in this early neonatal stage, or if their activity corresponds to other developmental functions.

Much of this lack of clarity has been due to the inability to isolate hunger and thirst in neonates as neonates satiate both drives by consuming maternal milk. In addition, given that the neonates cannot independently seek food or water, it also remains an open question if hunger or thirst alone would be sufficient to drive feeding. Here, we developed a continuous feeding platform for neonatal mice separate from the dam to isolate hunger and thirst response in the neonatal brain. We found that neural responses were already need-specific by the first postnatal week, well before the age at which mice can independently seek water or food. As in adult, ARC neurons responded specifically to hunger while SFO and OVLT neurons responded specifically to thirst in the neonate. Behaviorally, induced hunger or optogenetic activation of AgRP neurons drove feeding behaviors more effectively than induced thirst or activation of Agtr1a neurons in neonatal mice. *In vivo* recordings in neonates revealed that neurons in the dehydrated and starved neonatal mouse increased in activity in response to maternal presentation and that this increase was suppressed by feeding in the hypothalamus and thalamus, but not in the septal nuclei, hippocampus, or cortex. Interestingly, different subpopulations within MnPO, a region previously described in the thirst-responding circuit^19^, were active under hunger and thirst conditions in neonatal stages. Across ARC, SFO, OVLT, and MnPO, distinct neural representations of different need states were maintained in adulthood regardless of whether mice were introduced to food and water separately or if they were fed a liquid diet throughout their lifetime. Together, our findings demonstrate that hunger and thirst neurons become active in a need-specific manner before animals can independently seek food and water, and suggest that the specificity of their response properties arises innately.

## RESULTS

### Distinct neural responses to starvation and dehydration in neonatal mice

To isolate hunger and thirst drives in neonatal mice, we developed a continuous feeding platform to selectively induce hunger or thirst in neonatal mice (**Figure 1A, B**). Mice were raised in their homecage until P7–9, at which time they were separated from the dam and cannulated with a thin microurethane tube into the upper esophagus and isolated in a warmed chamber. Using this platform, mice could be fed with various liquids while isolated from the dam. Using a continuous pump to deliver these liquids, we found that feeding a mouse milk substitute^20^ 80–120 μl per hour for approximately 16 hours led to a similar weight gain as homecage mice who fed from their dam for a similar duration of time (**Figure 1C**). Thus, we used this milk substitute, feeding rate, and duration for the remainder of experiments using this feeding platform.

**Figure 1.**
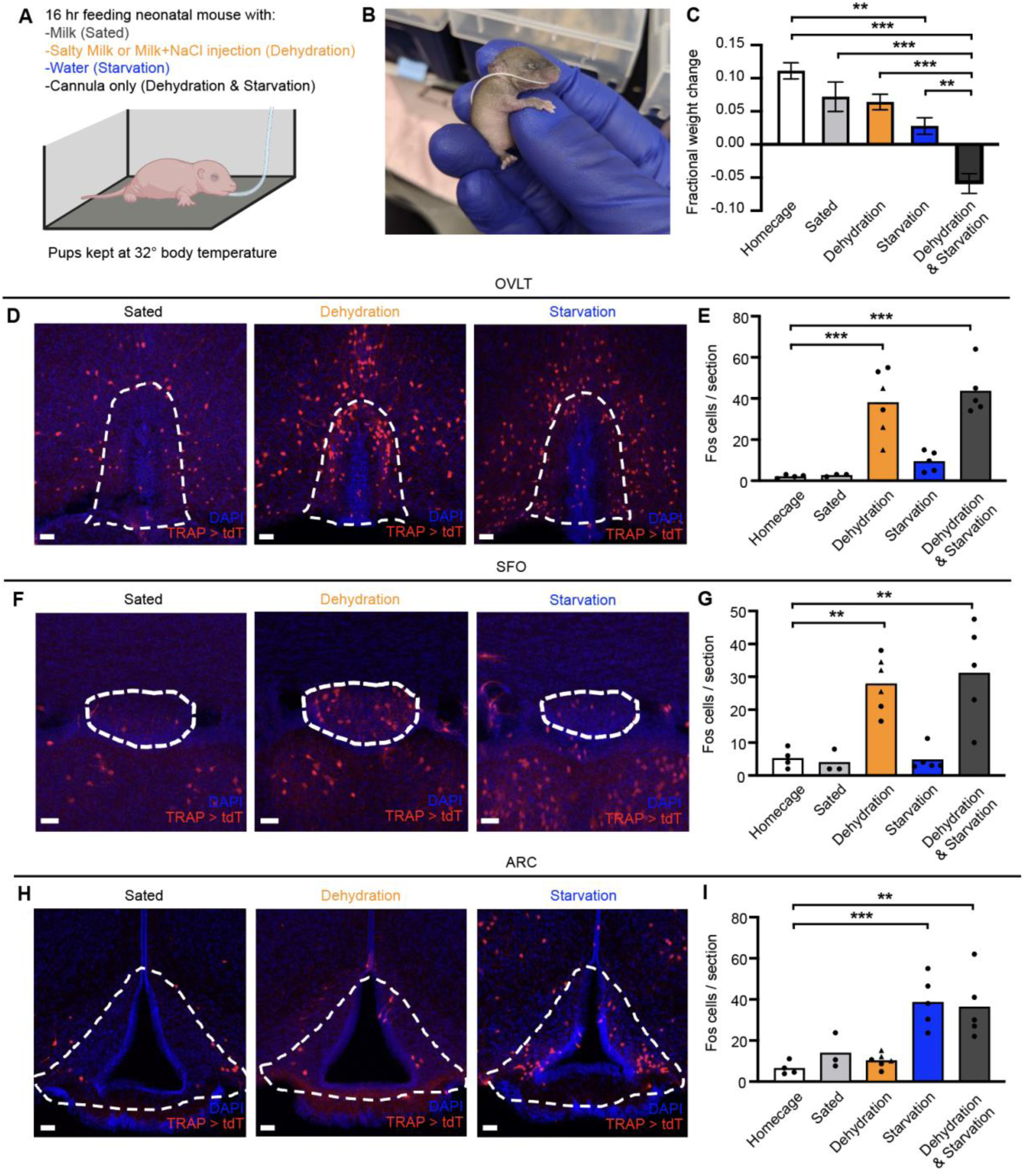
Distinct activation of hunger and thirst regions in neonatal mice by starvation and dehydration. (A) Schematic of neonatal mouse with an implanted esophageal cannula in a warmed chamber. (B) Photo of a cannulated P7 neonatal mouse. (C) Weight change of neonates after 16 hours of feeding in the cannula system, shown as fractional body weight change. n = 4–6 mice per condition, one-way ANOVA with post-hoc Tukey HSD. Error bars, SEM. (D, F, H) Example images of TRAPed cells in OVLT (D), SFO (F), and ARC (H) of neonatal mice in sated (fed milk), dehydration (fed salty milk, or fed milk with an NaCl injection 90 min prior to TRAPing), or starvation (fed water) conditions. Scale bar, 40 μm. (E, G, I) Quantification of TRAPed cells in OVLT (E), SFO (G), or ARC (I) in homecage, sated, dehydration, starvation, or dehydration & starvation (fed nothing) conditions. For dehydration conditions, circular points correspond to mice fed salty milk, and triangular points correspond to mice fed milk with an NaCl injection. n = 3–6 mice per condition, one-way ANOVA with post-hoc Tukey HSD with comparisons of all conditions to homecage condition. Abbreviations: OVLT, organum vasculosum laminae terminalis; SFO, subfornical organ; ARC, arcuate nucleus of the hypothalamus. For all figures, *p < 0.05, **p < 0.005, ***p < 0.001. See additional data in Figure S1.

To mimic a starvation condition, mice were fed water (distilled water with 35 mM NaCl, a hypotonic solution but reduces the chance of significant hyponatremia) for 16 hours before perfusion. To mimic a dehydration condition at the end of the experiment, mice were either fed salty milk for 16 hours (see **Methods**), or milk for 16 hours with 2M NaCl injection 90 min prior to TRAP labeling (see below). Starved and dehydrated mice were compared to “sated” mice that were fed milk and were also isolated in the warmed chamber for 16 hours to control for isolation from the dam and nest as a stimulus. In sated and dehydrated mice, feeding resulted in similar weight gains as homecage mice who were not removed from the dam and nest, whereas water feeding resulted in a smaller increase in weight likely due to lack of caloric intake ( **Figure 1C**). Mice that were cannulated but not fed with milk or water, mimicking a starvation-and-dehydration condition, showed a marked decrease in weight over the same period.

To examine the neuronal responses to starvation and dehydration in neonatal stages, we used the TRAP2^11,21^ mouse to facilitate permanent labeling of active neurons (determined by the expression of the activity-dependent immediate early gene *Fos*) in these different states. Using this approach, we found that early postnatal mice already demonstrate distinct neural responses to starvation and dehydration (**Figure 1D–I**). Starvation induced activity in ARC but not OVLT or SFO, whereas dehydration induced activity in OVLT and SFO but not ARC. These data demonstrate that hunger and thirst regions have distinct activation at this early postnatal stage. Interestingly, MnPO, a region that receives input from OVLT and SFO and has been implicated in the thirst but not hunger response^11,19^, did not have statistically significant differences between starvation and dehydration (**Figure S1A,B**). As will be described later, the neural responses induced by starvation and dehydration likely represent distinct subpopulations of neurons.

To confirm that starvation-activated neurons in ARC correspond to AgRP neurons, we combined *TRAP2* with *NPY-GFP* mice, as AgRP is known to be co-expressed with neuropeptide Y (NPY) in the ARC hunger neurons^2^. Here, we found that starvation-activated neurons labeled in P8 mice fed only water had high correspondence to NPY+ neurons when examined at P14–16 (**Figure S1C, D**). Co-staining with pro-opiomelanocortin (POMC), which is expressed in neurons that respond to satiation rather than starvation^3^, revealed low correspondence with TRAPed neurons in hungry mice at a similar age (**Figure S1C, D**). These findings suggest that the starvation-activated neurons in neonatal ARC largely represent AgRP hunger neurons.

Together, these findings suggest that the neural correlates of hunger and thirst are already present even before animals can perform distinct behaviors in response to starvation and dehydration.

### Neonatal feeding behaviors are driven more strongly by hunger than by thirst

Although neural responses to starvation and dehydration were already distinct in neonates, it is unclear the degree to which these responses would promote downstream feeding behavior. Thus, we next asked whether hunger or thirst states were sufficient to induce feeding behaviors. First, using our feeding platform, we induced hunger, thirst, or both in neonates (**Figure 2A**). We then assayed feeding motivation by measuring latency to attach to an anesthetized dam, total time attached to the dam, and weight change after free feeding. We compared these behaviors in dehydration and/or starved pups to those in sated pups (a pup isolated in the warmed chamber and fed milk). For behavioral experiments, thirst was only induced by feeding salty milk to remove the need to inject NaCl solutions shortly before the behavior session, which may cause disruptions in behavior, and to allow pups to have similar durations of induction of hunger and thirst.

**Figure 2.**
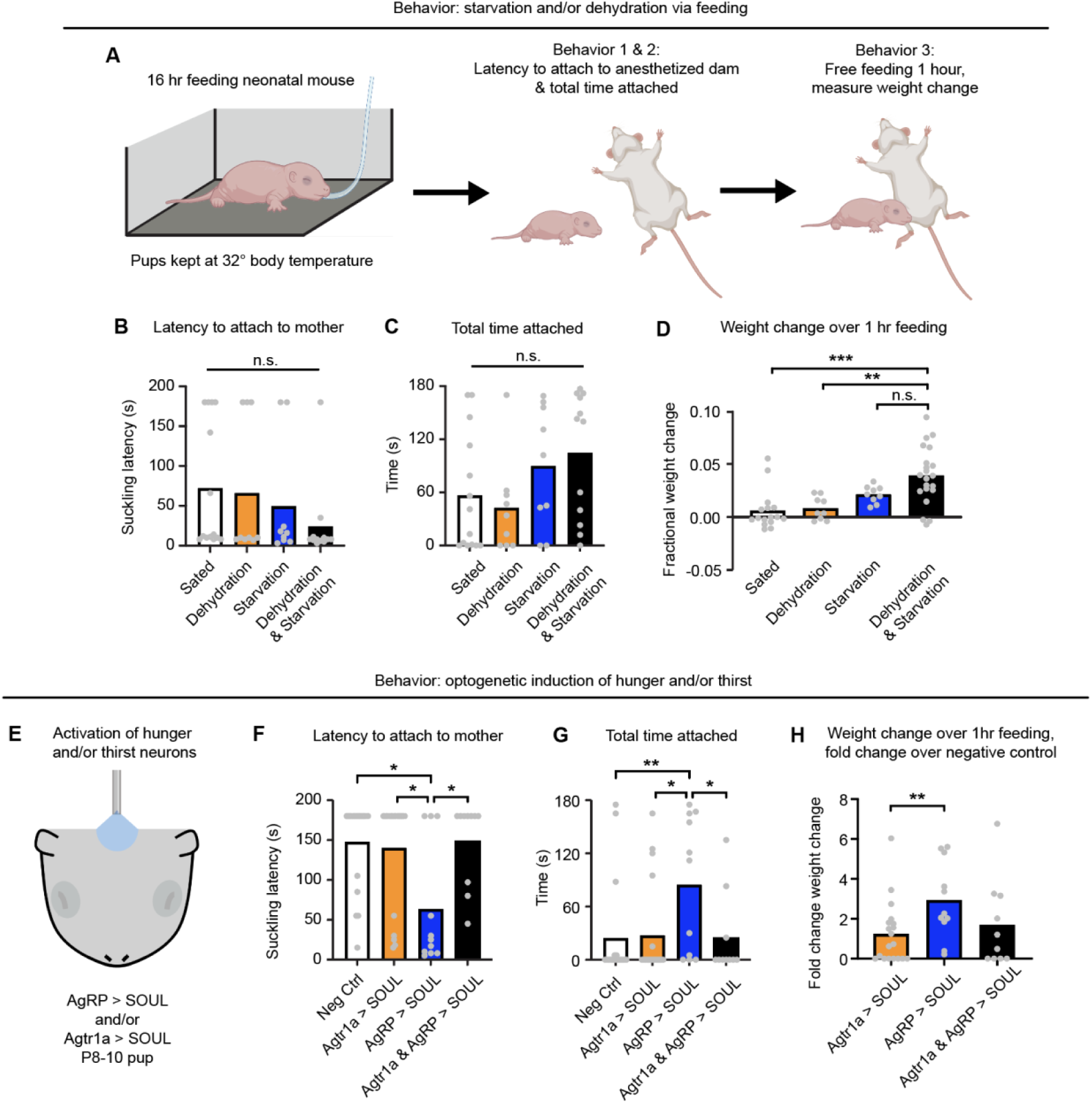
Attachment and feeding behaviors in thirst- and hunger-induced neonatal mice. (A) Schematic of attachment and feeding behaviors in a neonatal mouse after induction of hunger and/or thirst. (B) Quantification of latency to attach to mother after induction of hunger and/or thirst in neonatal mice via feeding. (C) Quantification of total time a neonate was attached to the mother for a 3-minute period after induction of hunger and/or thirst. (D) Quantification of fractional body weight change after a neonate was allowed to freely feed from the mother for 1 hour after induction of hunger and/or thirst. (E) Schematic showing transcranial activation of SOUL in *AgRP*- or *Agtr1a*-expressing neurons in a neonatal mouse. AgRP > SOUL, *AgRP-Cre/+;SOUL/+*. Agtr1a > SOUL, *Agtr1a- Cre/+;SOUL/+*. Agtr1a & AgRP > SOUL, *Agtr1a-*Cre/+;AgRP*-Cre/+*;*SOUL/+*. (F) Quantification of latency to attach to mother after induction of hunger and/or thirst in neonatal mice via SOUL. (G) Quantification of total time a neonate was attached to the mother for a 3-minute period after induction of hunger and/or thirst in neonatal mice via SOUL. (H) Quantification of fractional weight change after a neonate was allowed to freely feed from the mother for 1 hour after induction of hunger and/or thirst in neonatal mice via SOUL. For (C, D, G, H), n = 9–19 mice per condition, one-way ANOVA with post-hoc Tukey HSD. For (B, F), n = 9–19 mice per condition, Kruskal-Wallis test with post-hoc Dunn’s test. *p < 0.05, **p < 0.005, ***p <0.001. See additional data in Figure S2.

Following induction of starvation and/or dehydration, we found no significant difference in latency to attach or total time attached to the dam compared to the sated pup (**Figure 2B, C**). Interestingly, pup behaviors for latency to suckling demonstrated a bimodal distribution, where most mice attached immediately across all conditions. This may suggest that for suckling behavior, isolation from the mother may play a more significant role than hunger or thirst states. Alternatively, mice that were isolated for an extended period of time from the mother have been known to demonstrate reductions in maternal-seeking behaviors such as ultrasonic vocalizations, and thus a lack of a statistically significant difference here could be masked by this reduction^13^. It is possible that this occurred for the subset of mice which did not demonstrate any feeding behaviors. Regardless, these data suggest that attachment to the dam does not differ across hunger and thirst states when induced using our cannula feeding platform (**Figure 2A**, *behavior 1 & 2*).

Because suckling may be a readout of a bonding behavior in addition to hunger and thirst, we next used a more direct measure of consummatory behavior by assaying weight change over free feeding after starvation and/or dehydration states were induced (**Figure 2A**, *behavior 3*). We found that sated or dehydrated-only mice consumed significantly less milk than starved-and-dehydrated mice (**Figure 2D**). Starved-only mice, however, did not have significantly reduced milk consumption compared to starved-and-dehydrated mice (**Figure 2D**). This suggests that hunger drives feeding more similarly to a pup with both hunger and thirst. Interestingly, the fact that the magnitude of feeding was greater in starved-and-dehydrated mice than starved-only mice while dehydration only did not elicit much consumption suggests that hunger and thirst may still act synergistically. This was observed when we measured weight change in the initial phase of consumption, where starved-and-dehydrated mice did show a significant increase in weight gain over starved-only mice (**Figure S2C**). Together, these findings suggest that feeding may be driven more strongly by hunger than thirst in neonatal stages.

As an additional approach, we used the step-function opsin SOUL to activate AgRP and/or Agtr1a neurons to artificially induce a state of hunger and/or thirst, respectively^12,22^ (**Figure 2E**). Previous studies have shown that transcranial optogenetic stimulation in neonatal mice was sufficient to activate SOUL-expressing neurons, likely owing to the relative translucency of the skull at this early developmental stage^23^. We found that brief transcranial stimulation of neonates was indeed sufficient to activate the majority of AgRP or Agtr1a neurons expressing SOUL in ARC or SFO, respectively (**Figure S2A, B**). Thus, we used SOUL activation in these mice as a method to artificially induce hunger and thirst states in neonates without the need for extended isolation from the dam and nest.

Using this approach, we found that optogenetically inducing hunger but not thirst significantly increased attachment behaviors (**Figure 2F, G**). Interestingly, optogenetically inducing hunger and thirst simultaneously did not increase attachment behaviors, suggesting that while thirst alone does not drive these behaviors, activation of thirst neurons may disrupt hunger-driven behaviors as has been discussed in previous work on conflicts between internal states^24,25^. However, given Agtr1a neurons can be found in various regions^26^, we cannot rule out the possibility that our optogenetic manipulation activates with other neuronal circuits that interfere with these neonatal behaviors. Analysis of consummatory behaviors did not show a statistically significant increase in hunger-induced mice, whether after 1 hour of free feeding or in the initial phase of feeding (**Figure S2D, S2E**). However, this became statistically significant when normalizing for consummatory behavior in negative control littermates (**Figure 2H**). This normalization is an important step as initial satiation states between litters differ greatly, and brief optogenetic stimulation in otherwise sated pups is likely much more sensitive to these initial differences than induction of hunger or thirst through 16 hours of cannula feeding. Of note, mice across all conditions had weaker attachment and consummatory behaviors following optogenetic activation compared to those tested using our feeding platform, likely because these mice were not subjected to prolonged isolation which likely contributes to both attachment and consumption. However, the findings between our feeding platform and optogenetic stimulation consistently demonstrate a preferential increase in feeding behaviors in hunger states.

Taken together, these findings suggest that while isolation from the dam is likely a robust driver of feeding behaviors, dehydration and starvation further increase the drive for these behaviors. Hunger, whether induced by our feeding system or optogenetically, drives feeding behaviors more than does thirst in this early developmental stage.

### *In vivo* neuronal responses across various brain regions in the neonatal mouse to food and water

To study the neuronal responses in hunger and thirst states *in vivo*, we used Neuropixels to record the activity of neurons in starved-and-dehydrated neonatal mice in response to feeding of salty milk (to sate hunger but not thirst) or water (to sate thirst but not hunger) through an esophageal cannula (**Figure 3A**)^27^. Two probes were used simultaneously, to target SFO and ARC (**Figure S2A, B**). While we primarily targeted these nuclei, the large span of Neuropixels relative to the small neonatal brain meant that we also captured units in many regions across the brain during these recordings.

**Figure 3.**
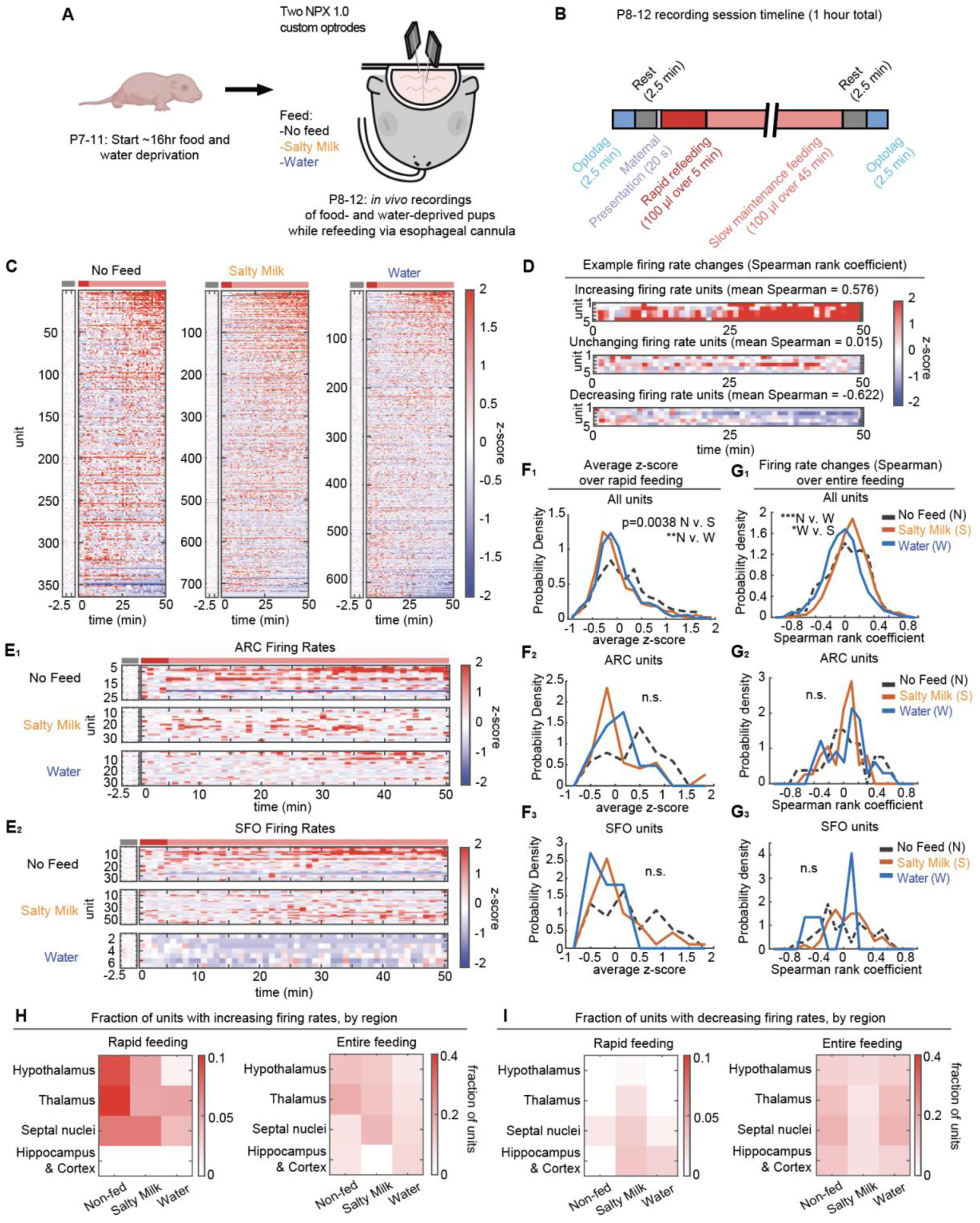
Distributed state-driven changes in neuronal activity when sating thirst or hunger *in vivo*. (A) Schematic of P3-TRAP of starvation- and dehydration-activated neurons, followed by dual Neuropixels optrode recordings in P8–12 mice that had been deprived of nourishment for ∼16 hours. Optotagging during probe insertion was used to aid targeting of SFO and ARC (see Methods). (B) Structure of 1-hour long recording session. (C) Firing rates of neurons across all units recorded during 50-minute refeeding of P8–12 mice not fed, fed salty milk (sating hunger), or fed water (sating thirst). Firing rates are shown as z-score averages in 1-min bins, relative to firing rates during the initial 2.5 min rest period as a baseline (n_No Feed_ = 364 units across 3 mice, n_Salty Milk_ = 733 units across 5 mice, n_Water_ = 631 units across 4 mice). (D) Firing rates of example units with increasing firing rates during refeeding (top), unchanging firing rates (middle), or decreasing firing rates (bottom), as measured by spearman rank coefficient of z-scored firing rates during 50-min refeeding. Mean spearman rank coefficients for these example units are shown. (E) Firing rates of units recorded during feeding of P8–12 mice not fed, fed salty milk (sating hunger), or fed water (sating thirst) for units in ARC (E_1_) or SFO (E_2_). (F) Distribution of firing rate changes of units in P8–12 mice across feeding conditions shown as average z-scored firing rates during rapid feeding for all units (F_1_), units in ARC (F_2_) or units in SFO (F_3_). (G) Distribution of firing rate changes of units in P8–12 mice across feeding shown as spearman rank coefficient of z-scored firing rates during 50-min feeding for all units (F_1_), units in ARC (F_2_), or units in SFO (F_3_). (H, I) Fraction of units within broad regions that have significant increases (H) or decreases (I) in firing rate (defined as a greater than one standard deviation change in average z-score during rapid feeding or spearman rank coefficient throughout feeding) across different feeding conditions. Hypothalamic units (n_No Feed_ = 166 units, n_Salty Milk_ = 188 units, n_Water_ = 104 units), thalamic units (n_No Feed_ = 62 units, n_Salty Milk_ = 253 units, n_Water_ = 279 units), septal nuclei units (n_No Feed_ = 124 units, n_Salty Milk_ = 262 units, n_Water_ = 176 units), and hippocampal & cortical units (n_No Feed_ = 12 units, n_Salty Milk_ = 30 units, n_Water_ = 72 units). For E–G, ARC units (n_No Feed_ = 25 units, n_Salty Milk_ = 36 units, n_Water_ = 31 units), SFO units (n_No Feed_ = 35 units, n_Salty Milk_ = 57 units, n_Water_ = 7 units). For all figures, one-way ANOVA with post-hoc Tukey HSD; *p < 3 x 10^-3^, **p < 3 x 10^-4^, ***p < 6 x 10^-5^; n.s. not significant, with Bonferroni correction for multiple comparisons. N v. W, No feed vs. Water, N v. S, No feed vs. Salty milk, W v. S, Water vs Salty milk. Gray bar indicates pre-feeding rest period, red bar indicates period of rapid refeeding (first 5 min of feeding), pink bar indicates slow maintenance refeeding (last 45 min of feeding). Abbreviations: ARC, arcuate nucleus; SFO, subfornical region. See additional data in Figures S3.

Prior to recording, we isolated P7–11 pups from the dam for 16 hours in a warmed chamber to induce both hunger and thirst (**Figure 3A**). We then implanted an esophageal cannula to feed the pup with salty milk or water to independently sate hunger or thirst, respectively. These were compared to mice in which a cannula was implanted but nothing was fed. Feeding occurred over 50 minutes, beginning with a 5 min rapid feed to quickly replenish food or water, followed by a slower maintenance feed over the next 45 min of recording (**Figure 3B**). Prior to feeding, we presented the maternal ventrum to the pups’ snout to study neuronal population responses in the dehydrated and starved state. During feeding, we measured changes in neuronal activity as z-scored firing rates relative to the baseline activity during the initial 2.5 min rest period (**Figure 3C–E**). We used two metrics to quantify the change in activity during feeding: the average z-score during the rapid feeding period to measure early responses (**Figure 3F**), and the Spearman rank coefficient over the entire feeding (a nonparametric measure of the general strength and direction of association between the activity and time) period to measure more gradual responses (**Figure 3G**). Spearman rank coefficients closer to 1 indicate consistent firing rate increases, whereas coefficients closer to –1 indicate consistent firing rate decreases, and coefficients close to 0 indicate little consistent change (**Figure 3D**).

We found that in units across brain regions, sating thirst leads to a decrease in neuronal activity during the rapid feeding period compared to no feeding (**Figure 3F_1_**) as well as continuous decrease over the 50-minute feeding period compared to no feeding or sating hunger (**Figure 3G_1_**). Sating hunger led to a qualitative decrease in activity during rapid feeding, but this did not reach significance when correcting for multiple comparisons (**Figure 3F_1_**, p = 0.0038). Sating hunger also did not lead to a decrease in population average activity over the 50-minute feeding period, perhaps in part due to the additional salt content increasing thirst and thus increasing activity a separate subpopulation of neurons (**Figure 3G_1_**). While our data were limited by a lack of access to specific cell types, we indeed found that broad brain regions contain both cells that significantly increase (**Figure 3H**) as well as those that significantly decrease (**Figure 3I**) their firing rates during feeding, suggestive of heterogeneous neuronal populations. Of note, we observed that only feeding salty milk led to a higher fraction of neurons significantly decreasing their activity during rapid feeding, potentially due to the caloric content present in salty milk but not water (**Figure 3I**).

Correspondingly, hypothalamic units demonstrated a significant decrease in firing during rapid feeding for both salty milk- and water-fed mice without consistent firing rate changes across the 50-minute feeding period (**Figure S3D–F**). The fact that rapid feeding decreased firing rates is consistent with previous findings that consumption rapidly decreased AgRP neuronal activity^28,29^. The lack of significant consistent firing rate changes across the 50 min feeding period may have contributions from satiety-related activity from populations such as POMC neurons, which is expected to increase in feeding but at a potentially slower timescale^3,28^. Units in the thalamus as well as septal nuclei also demonstrated changes in response to feeding ( **Figure S3D–F**), which may reflect state-driven changes in the integrative and gating functions in which these regions are implicated^30^. The septal nuclei have also been directly implicated in feeding and drinking behaviors^31,32^. In comparison, units within hippocampus and cortex did not have significant changes during any feeding period.

When comparing the firing rates within specific hunger- and thirst-related regions during feeding, we observe qualitative but not statistically significant decreases in activity during rapid feeding in ARC and SFO when sating hunger or thirst (**Figure 3E–G**). Units in MnPO also demonstrated the expected qualitative increases in activity with salty milk feeding and decreases in activity with water feeding, although no units were captured in this region in the non-fed condition (**Figure S3C**). The fact that rapid feeding of water led to a relative inhibition of some ARC units may be secondary to gut distension^33^, and rapid feeding of salty milk leading to a relative inhibition of some SFO units may be secondary to fluid-driven volume expansion^10^. Within ARC, SFO, and MnPO, lack of statistical significance despite observable qualitative differences are likely due to the relatively small number of isolated units within these regions. No differences in consistent changes in firing rates across the 50 min feeding period were observed in any of these regions, likely owing to our capturing of heterogeneous neuronal populations as previously discussed (**Figure 3G, S3C**).

Interestingly, we observed an overall increase in activity in the non-fed condition during the rapid feeding period (**Figure 3F_1_**, average z-score = 0.3363, p<0.0005, one sample t-test) which appears to be suppressed by feeding either salty milk or water. While this may be due to stress during a head-fixed recording session, such increased firing rate could also be in response to the pre-feed maternal presentation (**Figure 3B**). One possibility is that maternal presentation in a dehydrated and starved pup leads to activation of neural circuits involved in sensing the dam and driving feeding behavior, and that rapid feeding either salty milk or water suppresses this activation. Consistent with this interpretation, we observe similar suppression in the hypothalamus and thalamus (**Figure S3D–F**) as well as hunger- and thirst-related regions (**Figure 3E–G**) but not septal nuclei, hippocampus, or cortex.

In summary, feeding either salty milk or water leads to decreased activity in the hypothalamus and thalamus on shorter timescales, with more heterogeneous responses in activity on longer timescales. Increased activity in non-fed pups that is potentially due to presentation of maternal stimuli is suppressed when pups are fed with salty milk or water, which is consistent with our previous findings that feeding pups during isolation can reduce feeding behavior (**Figure 2**). Feeding salty milk leads to a qualitatively higher fraction of neurons with reductions in firing rates in the hypothalamus and thalamus on short timescales compared to the non-fed and feeding water conditions (**Figure 3H**, **3I**). Together, these findings demonstrate that *in vivo* firing rates of neurons across various brain regions are differentially sensitive to feeding in both dehydrated and starved neonates on different timescales.

### Neural representation of hunger and thirst is distinct by neonatal stages and does not depend on separation of food and water source

Given MnPO was activated in both hungry and thirsty neonatal mice (**Figure S1A, B**), we asked whether distinct subpopulations of neurons within MnPO respond to starvation and dehydration in neonatal mice. To accomplish this, we used TRAP2 to permanently label dehydration-activated (thirst-TRAPed) populations in MnPO in neonatal mice. 2–3 days following TRAPing, we induced starvation or dehydration and performed Fos staining to quantify the proportion of thirst-TRAPed neurons that were reactivated (**Figure 4A**).

**Figure 4.**
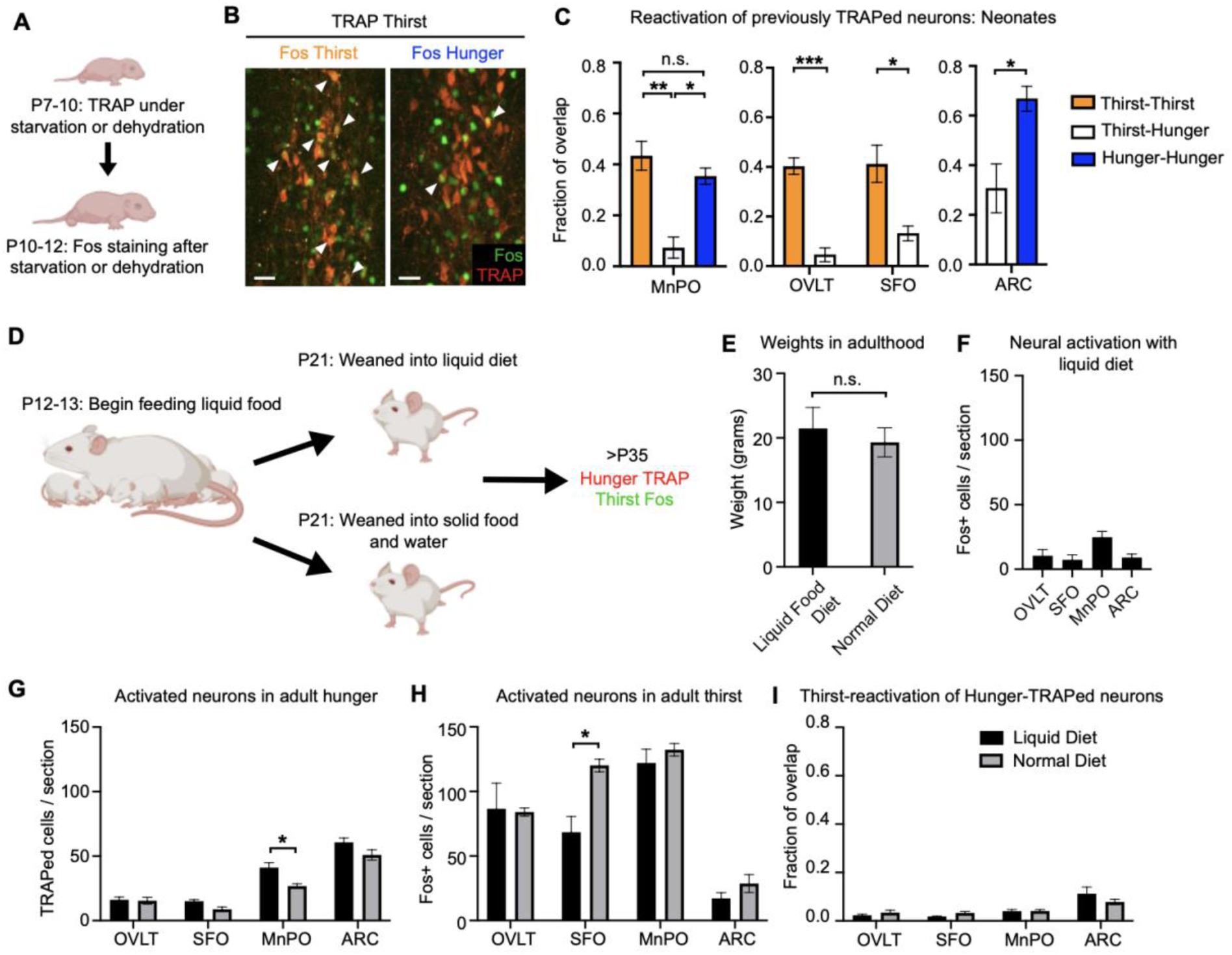
Starvation and dehydration activate distinct neuronal populations in neonates and adults fed a liquid diet. (A) Schematic showing thirst- or hunger-TRAP in a neonatal mouse followed by thirst- or hunger-Fos several days after TRAP. (B) Example image of Fos expression following thirst or hunger induction co-stained with thirst-TRAPed neurons, in neonatal MnPO. Arrowheads indicate double-labeled cells. Scale bar, 20 μm. (C) Quantification of Fos expression (indicating activation in thirst or hunger) in previously thirst- or hunger-TRAPed neurons in neonatal brain regions. Overlap is calculated as Fos+TRAP+ / TRAP+ neurons (n = 3–5 mice per condition. For OVLT, SFO, and ARC, unpaired t-test, * p < 0.05, *** p < 0.001. For MnPO, two-way ANOVA with post-hoc Tukey HSD; * p < 0.05, ** p < 0.005. Error bars, SEM. (D) Schematic of timeline for maintaining mice on a liquid diet. (E) Weights of P35 mice on a liquid diet and normal diet consisting of solid food and water (n = 5 mice per condition, unpaired t-test). Error bars, SEM. (F) Baseline Fos expression in hunger and thirst regions for mice on a liquid diet (n = 5 mice). (G, H) Quantification of activated neurons in adults fed a liquid diet or normal diet under induction of hunger (G) or thirst (H) (n = 5 mice per condition, two-way ANOVA with post-hoc Tukey HSD; * p < 0.05). Error bars, SEM. (I) Quantification of overlap of Fos expression (indicating activation in thirst) and TRAP labeling (indicating activation in hunger). Overlap is calculated as Fos+TRAP+ / Fos+ neurons (n = 5 mice per condition, two-way ANOVA with post-hoc Tukey HSD). Error bars, SEM.

We found significantly higher reactivation of thirst-TRAPed MnPO populations by dehydration than by starvation (**Figure 4B, C**). Similar results were found in thirst-TRAPed neurons in OVLT and SFO (**Figure 4C**). Likewise, we found significantly higher reactivation of hunger-TRAP neurons by starvation than by dehydration in MnPO and ARC (**Figure 4C**). Taken together, these data suggest that neuronal populations within hypothalamic regions responding to starvation and dehydration are already distinctly responsive to their given modality before mice can seek food and water separately.

Lastly, we asked whether these distinct neural responses in hunger and thirst could be altered by modifying the adult mouse diet such that food and water was obtained through the same source (**Figure 4D**). To do so, we created a liquid diet that was near-isotonic and nutritionally complete. Adult mice exclusively on the liquid diet were similar in weight to mice on normal diets (**Figure 4E**) and had very few Fos-expressing cells in the hunger and thirst regions, suggesting this diet was sufficient to sate both needs (**Figure 4F**).

We then examined neuronal activation patterns using hunger-TRAP followed by Fos staining under thirst condition in the same control mice or mice on liquid diet (**Figure 4D**). To induce hunger, we replaced liquid with water (for mice on the liquid diet) or removed food pellets (for control mice on normal diet consisting of solid food pellet and water) 36 hours prior to performing TRAP. Animals were subsequently returned to either liquid diet or normal diet. Later, thirst was induced by removing water (for control mice) or replacing liquid diet with solid food (for mice on the liquid diet) for 36 hours before perfusion and Fos staining. Mice on the liquid diet were also provided a small amount of salty liquid food (see Methods) to facilitate the switch in diet. As expected, hunger-TRAP predominantly captured neurons in ARC (**Figure 4G**), and thirst-Fos predominantly labeled neurons in OVLT and SFO (**Figure 4H**). TRAPed or Fos+ cells in hunger and thirst regions were similarly activated in hunger and thirst between mice on liquid and normal diet, with minor quantitative differences seen in the MnPO and SFO (**Figure 4G**, **4H**). Notably, mice in both conditions had little overlap between neurons activated under the hunger and thirst conditions (**Figure 4I**). In particular, MnPO, which had active neurons in response to both starvation and dehydration, had no increase in overlap in response to starvation and dehydration in mice that were only fed liquid diet prior to the final testing. This suggests that the distinction between neural correlates of hunger and thirst states does not depend on having the experience of separate food and water intakes in adult life.

## DISCUSSION

Proper neural responses to starvation and dehydration are vital for survival across the lifespan of an animal. Here, we develop a cannulation platform that allows prolonged feeding of neonatal mice isolated from the mother to distinguish the hunger and thirst states. Using this platform, we demonstrated that activation of hunger and thirst regions is specific to the respective needs in neonatal stages before the animal can separately seek food or water. Sated neonates, even when separated from the dam for prolonged periods of time, do not demonstrate increased maternal seeking behavior. Hunger states, whether induced optogenetically or starving but not dehydrating the animal, drives feeding behaviors more than does thirst in neonates. *In vivo* high-density single-unit recordings show that units across regions show relative reductions in activity within minutes of feeding, with activity changes at longer timescales becoming more heterogeneous. Lastly, in MnPO where cells are active in both hunger and thirst states, distinct subpopulations of neurons are activated in each state by early postnatal stages, and these distinct responses are not altered if an animal does not have separate access to food and water throughout their lifespan. These findings demonstrate that brain regions involved in responding to starvation and dehydration and regulating hunger- and thirst-driven behavior are established early and are robust to environmental changes later in life.

The early developmental specificity in neural correlates of hunger and thirst is significant given it is unclear at what stage these neurons develop the activation profiles seen in adults. For example, a previous study suggested that the activity of AgRP neurons in neonates correlate more to maternal interaction than starvation or feeding^13^. Although our experiments differed in the duration of starvation, maintenance of body temperature, and rate and continuity of feeding, our data demonstrates that at least a subset of neurons in ARC activated by prolonged starvation are suppressed by continuous feeding at this stage. Taken with our findings here, this may be consistent with the observation that AgRP neurons in adult mice respond to both nutritional status as well as food-predictive sensory inputs^29^. However, other studies have also shown that hunger and satiety hormones in development influence axonal growth of AgRP neurons rather than inducing changes in activity as seen in adult^16,17^. The switch to adult-like responses to hormones occurs around the time animals become independent and can seek food and water separately^18^. Axonal projections from AgRP neurons in ARC only reach a subset of their eventual target regions within the first postnatal week^15^, and thus early-maturing AgRP networks may be responsible for neonatally-relevant behaviors, while a fully mature AgRP network in the adult may be required to drive the complete repertoire of hunger-driven behaviors. Knowledge of when certain regions first become active under hunger and thirst conditions would thus help define critical windows during which alterations in hormonal signaling may affect maturation of such circuits. These possibilities highlight how future studies with more complete spatiotemporal map of hunger- and thirst-driven brain-wide activity across development would provide insight into the physiological maturation of circuits driving diverse behaviors across development, as well as how stage-dependent perturbations may lead to alterations at different connections within these networks.

Our novel platform for inducing starvation and dehydration as well as our use of SOUL in AgRP and Agtr1a neurons provides the first isolation of hunger and thirst states in neonates. While this facilitated the finding that hunger drives feeding behaviors more robustly than does thirst, there are three caveats to these findings. First, while feeding water through our cannulation system ensures starvation, injecting NaCl or feeding salty milk may not activate all thirst neurons given some thirst neurons respond to hypovolemia^10^. However, the substantial activation of thirst regions in our dehydration conditions suggests a thirst condition is likely induced. Second, SOUL activation in AgRP and Agtr1a neurons may drive downstream behaviors to different degrees. Even though SOUL activation leads to Fos expression in AgRP and Agtr1a neurons, Fos expression is a binary rather than continuous measure of activity. However, given that consummatory behaviors are not significantly different between starved mice and mice with both starvation and dehydration, we suggest that hunger does drive feeding more strongly than does thirst in neonates. Lastly, the consummatory behaviors we observe (i.e., weight gain with free feeding from the mother after induction of hunger and/or thirst) reflect the neonate’s drive to consume milk, which may differentially sate hunger and thirst. However, our finding that SOUL activation in AgRP neurons drive attachment behaviors more so than in Agtr1a neurons suggests that hunger does indeed drive feeding more strongly does than thirst in neonates.

The hunger- and thirst-responding regions examined here are important nodes for neural representation of homeostatic needs, but need-specific states often involve neuronal activity across many brain regions^11,25^. Here we provide the first multi-regional recordings demonstrating the neonatal neuronal response across different homeostatic states. In control non-fed mice, maternal presentation in the dehydrated and starved state led to a general increase in activity, while feeding either salty milk or water appeared to suppress this increase, particularly in hypothalamic regions of interest. This is consistent with our finding that sated pups do not demonstrate robust maternal seeking activity even when separated for prolonged periods of time. However, these neuronal populations do not demonstrate clear unidirectional activity changes in response to sating hunger and thirst on longer timescales. This is likely due to lack of targeting specific cell types in our recordings as we captured neurons with both significant increases and decreases in activity during a rapid response to consummatory cues and slower feeding to satiation. Future studies with cell-type-specific optotagging will be key to confirm this hypothesis and provide insight on how these different components of homeostatic circuits respond to feeding in starved and dehydrated states. Such studies would also facilitate investigation of cell types involved in more granular aspects of thirst or hunger that our feeding approach does not address, such as hypovolemic thirst or aspects of hunger related to stomach distension.

While one might expect circumventricular regions such as ARC, SFO, and OVLT to develop state-specific activation with starvation and dehydration early, it is surprising that downstream regions such as MnPO also have state-specific activation. This raises the question of what distinguishes these largely non-overlapping populations of dehydration- and starvation-activated neurons in MnPO. Recent work in thirst circuits of adult mice has identified transcriptomically distinct cell types that play different roles in the thirst-sensing process^9,10^. These studies show that different cell types respond to hypovolemic thirst, which promotes drinking to replenish fluid volume, and hyperosmotic thirst, which promotes drinking to normalize plasma osmolarity. Further work investigating when and how these cell types begin to separate into their distinct roles will provide more granular insight into the development of these otherwise early-established and robust circuits regulating hunger and thirst. For example, we found that mice on a liquid diet had decreased activation of SFO in response to dehydration and increased activation of MnPO in response to starvation. Reliance on a liquid diet likely alters baseline plasma osmolarity or volume, and investigations into which cell types may have decreased responses to dehydration will provide insight on how these circuits flexibly adapt to environmental changes.

## ACKNOWLEDGEMENTS

We thank Drs. Lisa Stowers, Darren Logan, Regina M. Sullivan, Kevin Franks, Lijun Qi, and Airi Yoshimoto as well as Jun H. Song for intellectual and technical guidance. We thank Dr. Guoping Feng for the SOUL Cre reporter mice. This work was supported by National Institutes of Health R01-NS050835 (L.L.). D.C.W. is supported by a Stanford Bio-X fellowship. L.L. is an HHMI investigator.

## AUTHOR CONTRIBUTIONS

D.C.W., Y.W., and L.L. designed the experiments. D.C.W., Y.W., and C.M. performed the experiments. D.C.W. and L.A.E. analyzed the data. D.C.W. collected the *in vivo* recording data with the help of F.S.-V., and K.F. D.C.W. and L.L wrote the manuscript with help from Y.W. and K.F. All the authors read and edited the final version of the manuscript.

## MATERIALS AND METHODS

### Animals

All animal procedures adhered to animal care guidelines from Stanford University’s Administrative Panel on Laboratory Animal Care.

To generate *TRAP2;Ai14* and *TRAP2;Ai32* mice, we crossed *Fos2A-iCreER* mice (FosTRAP2, Jackson Laboratory, Stock 030323)^21^ with tdTomato Cre reporter mice (*Ai14*, Jackson Laboratory, Stock 007914)^34^ or ChR2-eYFP Cre reporter mice (*Ai32*, Jackson Laboratory, Stock 012569)^34^. These mice were then backcrossed to wild-type CD1 mice twice, and bred to homozygosity for all alleles, resulting in *TRAP2/TRAP2;Ai14/Ai14* or *TRAP2/TRAP2;Ai32/Ai32* mice. Backcrossing to CD1 improved litter size and pup size to facilitate cannulation. To generate *AgRP-Cre;SOUL* and *Agtr1a-Cre;SOUL mice*, *SOUL-P2A-tdT* Cre reporter mice (*SOUL*, Jackson Laboratory 032301) were crossed to *AgRP-Cre* (AgRP, Jackson Laboratory 012899)^35^ or *Agtr1a-Cre* (*Agtr1a*, Jackson Laboratory 031487)^36^ mice. To generate *AgRP-Cre;Agtr1a-Cre;SOUL* mice, *AgRP-Cre;SOUL* mice were crossed to *Agtr1a-Cre;SOUL* mice. To generate *TRAP2/+;Ai14/+;NPY-hrGFP/+* mice, *TRAP2;Ai14* mice were crossed with *NPY-hrGFP* mice (NPY, Jackson Laboratory 006417)^37^.

### Tamoxifen preparation, administration, and neonatal mouse fostering post-administration

4-hydroxytamoxifen (4-OHT; Sigma H6278) at 20 mg/ml in ethanol was added to a 1:4 mixture of castor oil:sunflower seed oil (Sigma 259853 and Sigma S5007) to give a final concentration of 10 mg/ml 4-OHT. Ethanol was evaporated by vacuum centrifugation. Neonatal 4-OHT administration was 15mg/kg of animal body weight; adult 4-OHT administration was 50 mg/kg of animal body weight. All 4-OHT was administered intraperitoneally (IP).

### Histology

Animals were perfused transcardially with phosphate-buffered saline (PBS) followed by 4% paraformaldehyde (PFA). Brains were then post-fixed overnight at 4°C in 4% PFA. Coronal sections were then cut at 60-µm thickness using a Leica Vibratome system. Immunostaining was performed with primary antibodies for c-Fos (Synaptic Systems 226 008), tdTomato (Rockland 600-401-379), GFP (Abcam Ab13970), or POMC (Phoenix Pharmaceuticals 27-52) in PBST (PBS + 0.1% Triton) at 4°C for two nights, washed 3 x 10 min with PBST, secondary antibodies in PBST overnight at room temperature, washed 3 x 10 min with PBST, and mounted in Vectashield (Vectorlabs).

### Isolation of hunger and/or thirst states in neonatal mice

To selectively induce hunger or thirst states in neonatal mice, esophageal cannulation was performed to provide continuous feeding of liquids. Mice were removed from the homecage, weighed, and briefly anesthetized with 3% isofluorane to facilitate implanting the feeding tube.

The feeding tube, a 0.025” OD polyurethane tubing (Braintree MRE025), was attached to a 27G needle on a 3 ml syringe containing milk (Organic Valley Half & Half), salty milk (half & half with solid NaCl added to a final concentration of 100 mM), or water (with 35 mM NaCl). The end of the tube was briefly heated over flame to create a curve in the tubing. The curved tip of the tube was dipped in milk and inserted into the pup’s oral cavity until the end of the tube just entered the esophagus (a slight increase in insertion resistance occurred when the tube hit the epiglottis, and entrance into the esophagus provided relief of that resistance). Shallow esophageal cannulation (located to upper esophagus) was necessary for feeding over many hours, as mice at this age do not reflexively ingest liquid directly placed in the oral cavity, and cannulation deeper than this compresses thoracic structures leading to respiratory difficulty and high rates of mortality.

After cannulation, a small dab of superglue was applied to adhere the curved part of the tubing to the cheek of the pup, and the pup was allowed to wake from isoflurane anesthesia. Pups were then placed in isolated chambers on a heat pad and kept at approximately 32°C. Milk, salty milk, or water was then pumped at a rate of 80-120 μl per hour (New Era Pump Systems NE-1200) for approximately 16 hours before 4-OHT administration for TRAP, or perfusion for Fos staining.

To induce both hunger and thirst in neonatal mouse mice, pups taken from homecage were cannulated as described above and kept at 30-32°C in an isolated chamber for approximately 16 hours but without any milk or water infusion prior to TRAPing or perfusion.

To induce thirst in neonatal mouse mice, fully sated mice received IP injections of 2 M NaCl (5 ul / g bodyweight). For TRAP, 15 mg/kg 4-OHT was administered 90 min after NaCl injection, and mice were returned to their dam 3 hours afterwards. For experiments in which previously TRAPed neurons were compared to Fos staining in P10-12 mice, thirst was induced by feeding milk followed by 2 M NaCl injection 60 min prior to perfusion for Fos staining.

### Feeding behavioral experiments in neonates

Dams were anesthetized with IP injections of ketamine (100 mg/kg) and xylazine (10 mg/kg), followed by IP injection of oxytocin (4 IU/kg) to facilitate milk letdown. Prior to each experiment, milk letdown was manually confirmed. Each litter used within a behavioral session contained mice that were sated, hungry, thirsty, or both, typically 1–2 mice per condition.

Anesthetized dams were placed supine in a small chamber over nesting and homecage bedding, and mice were placed approximately 2 cm from a nipple on the dam’s ventrum. Mice were not manually held during behavioral assays. Latency to attach was measured as the time elapsed before the pup began suckling from a nipple. Mice were then observed for 3 min, and the total time attached within these 3 min was recorded. Mice who did not attach were given a latency of 3 min. Mice were then removed from the mother, and the nipple was covered with tape before placing the next pup 2 cm in front of a different nipple.

After attachment behaviors were completed for each pup, dams received a second dose of ketamine, xylazine, and oxytocin. All mice were then returned to the anesthetized dam by placing them directly on top of a nipple (to promote attachment and feeding), and allowed to freely feed for 1 hour. Pup weights were recorded before feeding, 3 minutes after feeding during the suckling behaviors, after 1 hour of free feeding. Mice that were stressed or demonstrated impaired motility during behavior were removed from analysis.

### SOUL activation in neonatal mice

P8–10 mice were isolated with their littermates from dam for approximately 45 minutes prior to behavioral assay. SOUL was then activated in AgRP > SOUL and/or Agtr1a > SOUL mice via transcranial stimulation for 30 s using 20-mW 488-nm laser by placing a bare optic fiber over the skin covering over lambda. Transcranial optogenetic stimulation has been previously demonstrated to be sufficient to activate SOUL-expressing neurons^23^. No surgery was performed on these animals. Following stimulation, mice were allowed to rest with littermates isolated from dam for approximately 5 minutes prior to behavioral assays. Behavioral assays were then executed as described above.

### Liquid diet composition and delivery

To maintain animals on a completely liquid diet, Ensure Complete Nutrition was mixed with whole milk at a 1:2.5 ratio. This yields a solution of approximately 350 mOsm, similar to the osmolality of many electrolyte replacement solutions. A plastic container containing 100 ml of this liquid mixture was placed in the mouse cage and replaced daily. Feeding of liquid began at P13–14 (soon after eye opening and before mice begin to explore food and water sources outside of maternal milk). Litters were weaned at P21, and half the litter continued on this liquid diet while the other half of the litter was given access to water and solid food pellets. Liquid diets were continued until mice were >P35 at which point the mice were used in experiments.

### Isolation of hunger or thirst states in adult mice

To induce hunger in adult mice, food was removed from the cage for 36 hrs, and mice were given *ad libitum* access to regular water. To induce thirst in adult mice, water was removed from the cage for 36 hrs, and mice were given ad libitum access to regular food pellets. Mice previously on a liquid-only diet were also given 2 ml / mouse of the liquid (Ensure + whole milk) with 500 mM NaCl at the beginning of the thirst-induction period to facilitate the switch to solid food. 50 mg/kg 4-OHT was then administered for TRAPing. Food or water was returned 6–12 hrs later, or animals were perfused for Fos staining.

### Neuropixels recording

Mice were isolated in an approximately 30-32°C chamber without feeding for approximately 16 hours prior to recording to induce hunger and thirst. Neonatal mice were awake and unanesthetized for the duration of the recording. Mice were headfixed and placed on a custom 3-D printed chamber to allow for a more natural head angle during recording and kept at approximately 30-32°C with a heatpad. Each mouse was only recorded in one session. After recording, mice were perfused for histological analysis to match probe depth to brain region. Region mapping was done manually using the Allen Brain Atlas^38^. Each mouse had two electrodes implanted simultaneously.

*In vivo* electrophysiological recordings of awake P8–12 mice were performed using Phase 3B Neuropixels electrodes with 384 active recording sites along the tip of the probe. Kwikcast above the craniotomy was removed and the craniotomy was filled with 0.9% NaCl in water. Electrodes were coated with a red lipophilic dye (DiI, Thermo Fisher) and dried for 3–5 min. Electrodes were then lowered slowly (2–3 μm/s) into brain using a micromanipulator (Scientifica PatchStar) to the desired depth or until the electrode bent, indicating that it had reached the bottom of the skull, after which the electrode were retracted 100 µm. Electrodes were then allowed to settle in the brain for 45 min before recording. Recordings were sampled at 30 kHz amplified (500x gain) and bandpass filtered (0.3–10 kHz). Signals were digitized with a CMOS amplifier and multiplexer on the Neuropixels electrode, then written to the disk using SpikeGLX software. Respiration was measured with a microbridge mass airflow sensor (Honeywell AWM3300V) positioned directly opposite the animal’s nose and signals were recorded using a MCC DAQ board. Neuropixels and MCC data were synchronized offline.

To optotag TRAPed units in *TRAP2/TRAP2;Ai32/Ai32* mice, we attached a 400-µm 0.39-NA optic fiber (Thor Labs) to each Neuropixels probe, approximately 500 µm lateral to the probe with the fiber tip terminating ∼1.5 mm from the tip of the electrode. A 470-nm LED (∼15 mW at the fiber tip). These were implanted into *TRAP2/TRAP2;Ai32/Ai32* mice that had 15mg/kg 4-OHT injected at P3 after 16 hours of food and water deprivation in order to express ChR2 in neurons active during starvation and dehydration. Given the regions of interest (SFO and ARC) are small and difficult to target, we recorded multiunit responses to 1 s 0.25 Hz light pulses presented through this fiber during probe insertion to guide probe targeting to starvation- and dehydration-responsive regions such as SFO and ARC. During the recording session, we presented brief pulses at the beginning and end of each recording (30 pulses of 20 ms at 0.5 Hz, and 5-10 pulses of 1 s at 0.25 Hz), the responses of which were later used to attempt to determine optotagged units. While we were unable to ultimately optotag sufficient numbers of sorted single units, the 1 s pulses during probe insertion increased our rate of successfully targeting SFO and ARC.

During the recording session, mice were implanted with esophageal cannula as described above. Optotagging was done within the first 2.5 minutes of the recording session. Mice were then allowed to rest for 2.5 minutes. The maternal ventrum was manually presented 2 cm from the pup snout for 20 sec after this rest period. Subsequently, mice were fed 100 µl of salty milk (milk with 100 mM NaCl) or water over 5 minutes via cannula. Mice were then fed another 100 µl of the same solution over the next 45 minutes. At the end of this 50 minutes of feeding, mice had a final 2.5 minutes of optotagging prior to conclusion of the recording session.

### Neuropixels spike sorting and analysis

Spike sorting was performed offline using Kilosort2.5 with default parameters. The clusters were manually curated using Phy2 to determine whether they were “Good” (well-isolated and stable) or “MUA” (poorly isolated or unstable units). All units classified by Kilosort as “Good” were manually inspected, and clusters with high signal-to-noise ratio, clean spike time auto-correlograms were kept for analysis.

A custom MATLAB script was written to analyze firing rates across recording session. Firing rates were calculated over 1 sec time bins, and a z-score for each time bin was calculated relative to the average firing rate across the recording. Activity was then smoothed using a moving average filter of 5 sec time bins. For analysis of firing rate across a 50 min recording session, z-scored activity was plotted in 1 min time bins calculated from the average of z-score within each minute relative to the baseline firing rate during the initial 2.5 min rest period prior to feeding. Change in z-score was calculated in MATLAB as the either the average z-scored firing rate within the rapid feeding period or Spearman rank coefficient of z-score in 1-min bins across the 50 min feeding period. Spearman rank coefficients were not used during the rapid feeding as temporal resolution of solution reaching the stomach was poor due to limitations of the pump and minor variations in esophageal cannulation depth. Spearman rank coefficients also perform poorly for non-monotonic time series, limiting their use to activity changes on shorter timescales, such as a unit decreasing activity rapidly followed by a slower increase in activity back to baseline.

---

**Figure S1.**
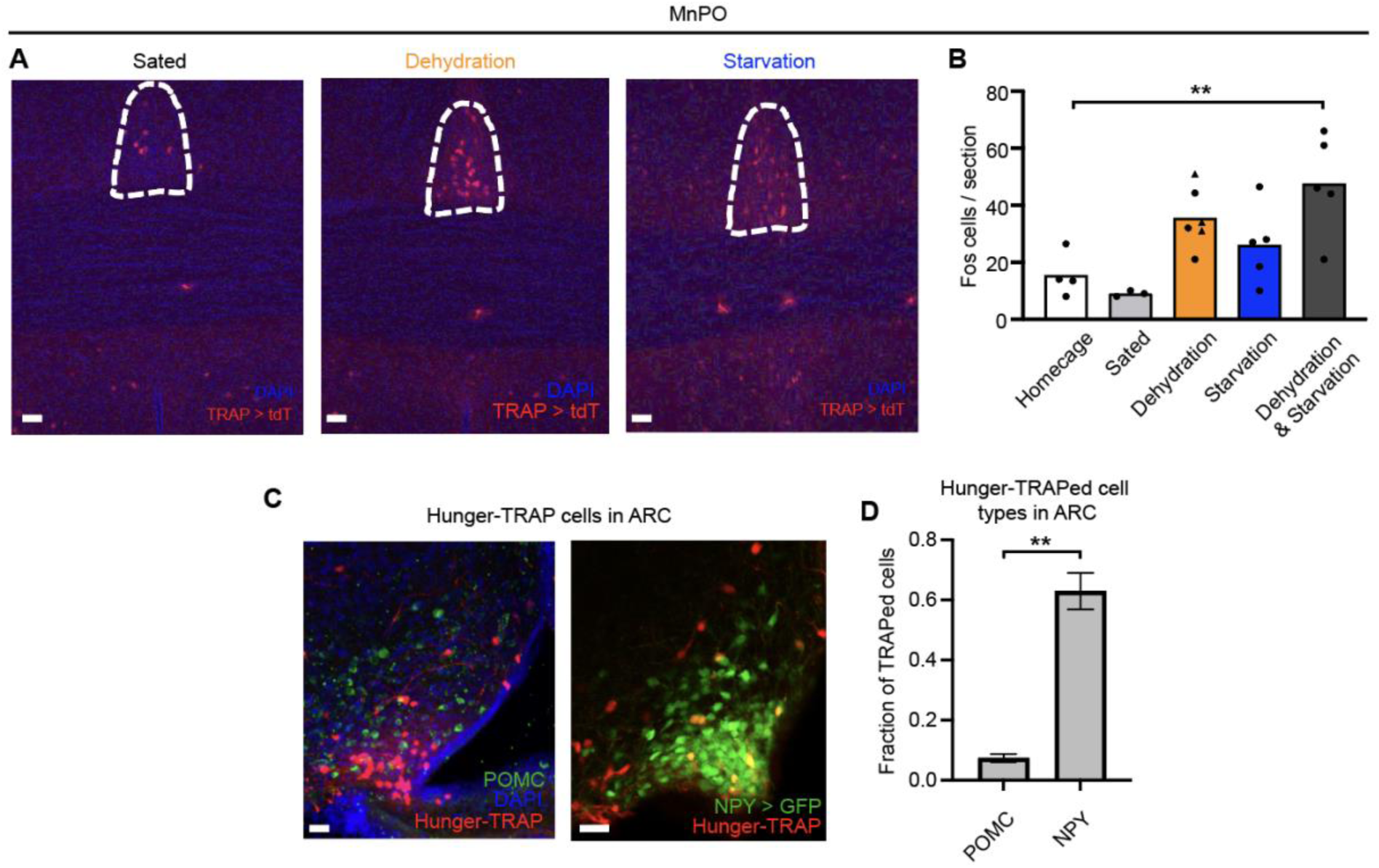
Hunger- and thirst-TRAP in MnPO, and identity of hunger-TRAPed cells, related to Figure 1. (A) Example images of TRAPed cells in MnPO of neonatal mice in sated (fed milk), dehydration (fed salty milk, or fed milk with an NaCl injection 90 min prior to TRAPing), or starvation (fed water) conditions. Scale bar, 50 μm. (B) Quantification of TRAPed cells in MnPO of homecage, sated, dehydration, starvation, or dehydration & starvation (fed nothing) conditions. For dehydration conditions, circular points correspond to mice fed salty milk, and triangular points correspond to mice fed milk with an NaCl injection. n = 3–6 mice per condition, one-way ANOVA with post-hoc Tukey HSD with comparisons of all conditions to homecage condition. **p < 0.005. (C) Example image showing overlap of P8 hunger-TRAPed cells with POMC (left) or NPY (right) stained at P14–16 in sections of the arcuate nucleus (ARC); NPY and AgRP are normally co-expressed in the ARC. (D) Quantification of fraction of P8 hunger-TRAPed cells that express POMC or NPY at P14–16. n = 3 mice per condition. Unpaired t-test; **p < 0.005. Error bar, SEM. Abbreviations: MnPO, median preoptic nucleus; ARC, arcuate nucleus of the hypothalamus.

**Figure S2.**
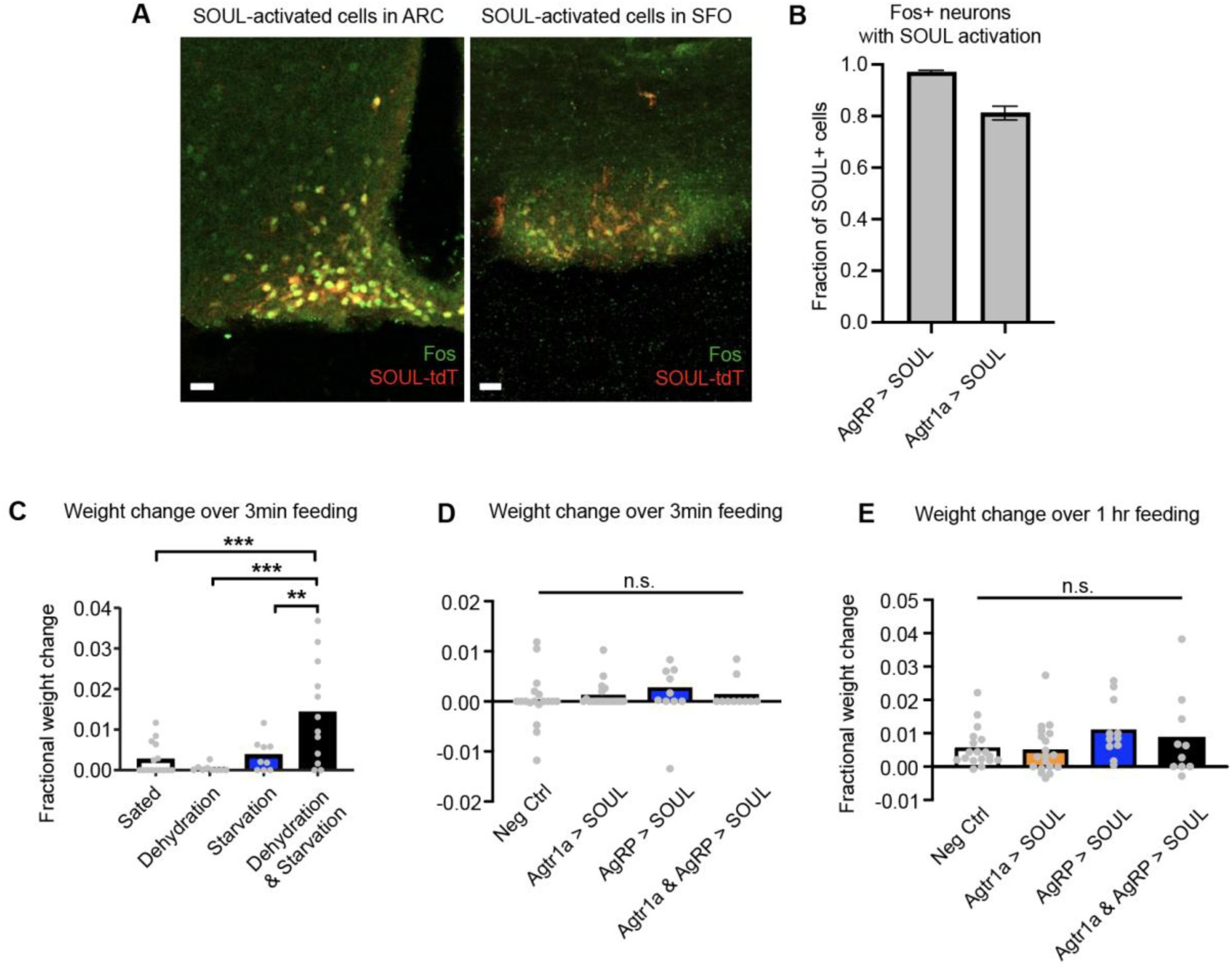
Activation of *AgRP*- and *Agtr1a*-expressing cells with SOUL, and additional feeding behaviors in thirst- and hunger-induced neonatal mice, related to Figure 2. (A) Example image showing Fos expression in AgRP > SOUL mice (left) and Agtr1a > SOUL mice (right) after transcranial activation of SOUL. Scale bar, 30 μm. (B) Quantification of fraction of SOUL+ neurons that express Fos after activation of SOUL. n = 3 mice per condition. Error bars, SEM. (C) Quantification of fractional body weight change within the first 3 minutes after a neonate was allowed to freely feed from the mother after induction of hunger and/or thirst via feeding milk (sated), salty milk (dehydration), water (starvation), or nothing (dehydration & starvation). (D) Quantification of fractional weight change within the first 3 minutes after a neonate was allowed to freely feed from the mother after induction of hunger or thirst in neonatal mice via SOUL. (E) Quantification of fractional weight change after a neonate was allowed to freely feed from the mother for 1 hr after induction of hunger or thirst in neonatal mice via SOUL, with fractional weight changes normalized by fractional weight changes of negative control littermates. For (C–E), n = 9–19 mice per condition, one-way ANOVA with post-hoc Tukey HSD. *p < 0.05, **p < 0.005, ***p < 0.001.

**Figure S3.**
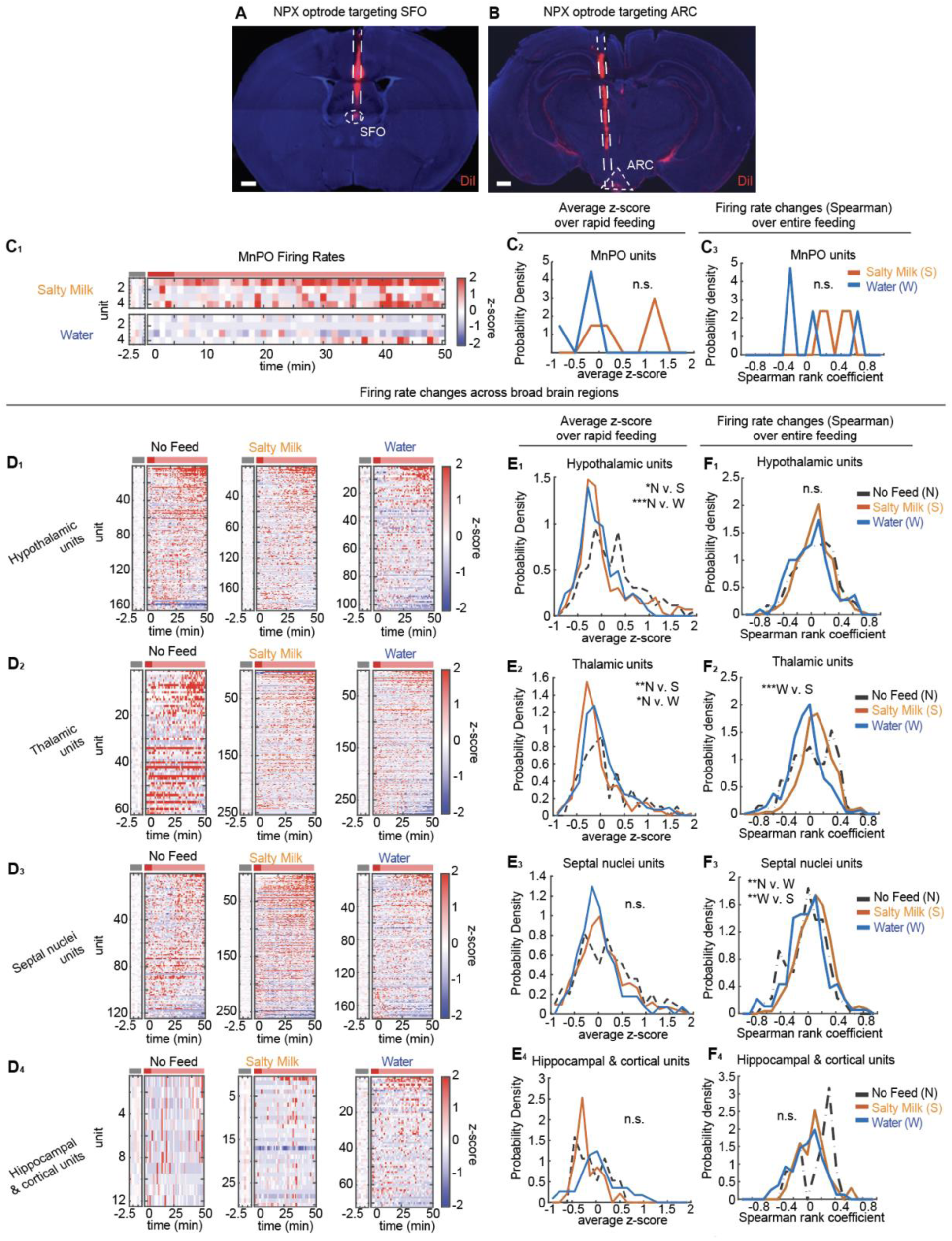
Characterization of region-specific state-driven changes in neuronal activity when sating thirst or hunger *in vivo*, related to Figure 3. (A, B) Schematic of Neuropixels track through the brain during dual optrode recordings, with one optrode targeting SFO (A) and one optrode targeting ARC (B). Scale bar, 500 μm. (C) Firing rates of units recorded during feeding of P8–12 mice fed salty milk (sating hunger), or fed water (sating thirst) for units in MnPO shown as z-scored firing rates across entire feeding (C_1_), average z-scored firing rates during rapid feeding (C_2_), and spearman rank coefficient of z-scored firing rates during 50-min feeding (C_3_). (D) Firing rates of units recorded during feeding of P8–12 mice not fed, fed salty milk (sating hunger), or fed water (sating thirst) for units in hypothalamus (D_1_), thalamus (D_2_), and septal nuclei (D_3_), and hippocampus & cortex (D_4_). (E) Distribution of firing rate changes of units in P8–12 mice across feeding conditions shown as average z-scored firing rates during rapid feeding for units in hypothalamus (E_1_), thalamus (E_2_), and septal nuclei (E_3_), and hippocampus & cortex (E_4_). (F) Distribution of firing rate changes of units in P8–12 mice across feeding conditions shown as spearman rank coefficient of z-scored firing rates during 50-min feeding units in hypothalamus (F_1_), thalamus (F_2_), and septal nuclei (F_3_), and hippocampus & cortex (F_4_). For all figures, hypothalamic units (n_No Feed_ = 166 units, n_Salty Milk_ = 188 units, n_Water_ = 104 units), thalamic units (n_No Feed_ = 62 units, n_Salty Milk_ = 253 units, n_Water_ = 279 units), septal nuclei units (n_No Feed_ = 124 units, n_Salty Milk_ = 262 units, n_Water_ = 176 units), and hippocampal & cortical units (n_No Feed_ = 12 units, n_Salty Milk_ = 30 units, n_Water_ = 72 units). For all figures, firing rates are shown as z-score averaged over 1-min bins relative to the initial 2.5 min rest period as a baseline. Gray bar indicates pre-feeding rest period, red bar indicates period of rapid refeeding (first 5 min of feeding), pink bar indicates slow maintenance refeeding (last 45 min of feeding). One-way ANOVA with post-hoc Tukey HSD; *p < 3 x 10^-3^, **p < 3 x 10^-4^, ***p < 6 x 10^-5^; n.s. not significant, with Bonferroni correction for multiple comparisons. N v. W, No feed vs. Water, N v. S, No feed vs. Salty milk, W v. S, Water vs Salty milk. Abbreviations: ARC, arcuate nucleus; SFO, subfornical region; MnPO, median preoptic nucleus.

## REFERENCES

1. Augustine, V., Lee, S. & Oka, Y. Neural Control and Modulation of Thirst, Sodium Appetite, and Hunger. Cell 180, 25–32 (2020).

2. Luquet, S., Perez, F. A., Hnasko, T. S. & Palmiter, R. D. NPY/AgRP Neurons Are Essential for Feeding in Adult Mice but Can Be Ablated in Neonates. Science 310, 683–685 (2005).

3. Cone, R. D. Anatomy and regulation of the central melanocortin system. Nat Neurosci 8, 571–578 (2005).

4. Aponte, Y., Atasoy, D. & Sternson, S. M. AGRP neurons are sufficient to orchestrate feeding behavior rapidly and without training. Nat Neurosci 14, 351–355 (2011).

5. Atasoy, D., Betley, J. N., Su, H. H. & Sternson, S. M. Deconstruction of a neural circuit for hunger. Nature 488, 172–177 (2012).

6. Betley, J. N., Cao, Z. F. H., Ritola, K. D. & Sternson, S. M. Parallel, Redundant Circuit Organization for Homeostatic Control of Feeding Behavior. Cell 155, 1337–1350 (2013).

7. Wu, Q., Boyle, M. P. & Palmiter, R. D. Loss of GABAergic Signaling by AgRP Neurons to the Parabrachial Nucleus Leads to Starvation. Cell 137, 1225–1234 (2009).

8. Dietrich, M. O., Zimmer, M. R., Bober, J. & Horvath, T. L. Hypothalamic Agrp Neurons Drive Stereotypic Behaviors beyond Feeding. Cell 160, 1222–1232 (2015).

9. Oka, Y., Ye, M. & Zuker, C. S. Thirst driving and suppressing signals encoded by distinct neural populations in the brain. Nature 520, 349–352 (2015).

10. Pool, A.-H. et al. The cellular basis of distinct thirst modalities. Nature 588, 112–117 (2020).

11. Allen, W. E. et al. Thirst-associated preoptic neurons encode an aversive motivational drive. Science 357, 1149–1155 (2017).

12. Leib, D. E. et al. The Forebrain Thirst Circuit Drives Drinking through Negative Reinforcement. Neuron 96, 1272–1281.e4 (2017).

13. Zimmer, M. R., Fonseca, A. H. O., Iyilikci, O., Pra, R. D. & Dietrich, M. O. Functional Ontogeny of Hypothalamic Agrp Neurons in Neonatal Mouse Behaviors. Cell 178, 44–59.e7 (2019).

14. Iyilikci, O., Zimmer, M. R. & Dietrich, M. O. Development of “Hunger Neurons” and the Unanticipated Relationship Between Energy Metabolism and Mother-Infant Interactions. Biological Psychiatry 91, 907–914 (2022).

15. Bouret, S. G., Draper, S. J. & Simerly, R. B. Formation of Projection Pathways from the Arcuate Nucleus of the Hypothalamus to Hypothalamic Regions Implicated in the Neural Control of Feeding Behavior in Mice. J. Neurosci. 24, 2797–2805 (2004).

16. Bouret, S. G., Draper, S. J. & Simerly, R. B. Trophic Action of Leptin on Hypothalamic Neurons That Regulate Feeding. Science 304, 108–110 (2004).

17. Steculorum, S. M. et al. Neonatal ghrelin programs development of hypothalamic feeding circuits. J. Clin. Invest. 125, 846–858 (2015).

18. Baquero, A. F. et al. Developmental Switch of Leptin Signaling in Arcuate Nucleus Neurons. Journal of Neuroscience 34, 9982–9994 (2014).

19. McKinley, M. J. et al. The median preoptic nucleus: front and centre for the regulation of body fluid, sodium, temperature, sleep and cardiovascular homeostasis. Acta Physiol 214, 8–32 (2015).

20. Görs, S., Kucia, M., Langhammer, M., Junghans, P. & Metges, C. C. Technical note: Milk composition in mice—Methodological aspects and effects of mouse strain and lactation day. Journal of Dairy Science 92, 632–637 (2009).

21. DeNardo, L. A. et al. Temporal evolution of cortical ensembles promoting remote memory retrieval. Nat Neurosci 22, 460–469 (2019).

22. Gong, X. et al. An Ultra-Sensitive Step-Function Opsin for Minimally Invasive Optogenetic Stimulation in Mice and Macaques. Neuron 107, 38–51.e8 (2020).

23. Wang, D. C., Santos-Valencia, F., Song, J. H., Franks, K. M. & Luo, L. Embryonically active piriform cortex neurons promote intracortical recurrent connectivity during development. Neuron 112, 2938–2954.e6 (2024).

24. Richman, E. B., Ticea, N., Allen, W. E., Deisseroth, K. & Luo, L. Neural landscape diffusion resolves conflicts between needs across time. Nature 623, 571–579 (2023).

25. Eiselt, A.-K. et al. Hunger or thirst state uncertainty is resolved by outcome evaluation in medial prefrontal cortex to guide decision-making. Nat Neurosci 24, 907–912 (2021).

26. Gonzalez, A. D. et al. Distribution of angiotensin type 1a receptor-containing cells in the brains of bacterial artificial chromosome transgenic mice. Neuroscience 226, 489–509 (2012).

27. Jun, J. J. et al. Fully integrated silicon probes for high-density recording of neural activity. Nature 551, 232–236 (2017).

28. Mandelblat-Cerf, Y. et al. Arcuate hypothalamic AgRP and putative POMC neurons show opposite changes in spiking across multiple timescales. eLife 4, e07122 (2015).

29. Chen, Y., Lin, Y.-C., Kuo, T.-W. & Knight, Z. A. Sensory Detection of Food Rapidly Modulates Arcuate Feeding Circuits. Cell 160, 829–841 (2015).

30. Risold, P. Y. & Swanson, L. W. Connections of the rat lateral septal complex 1Published on the World Wide Web on 2 June 1997. 1. Brain Research Reviews 24, 115–195 (1997).

31. Lorens, S. A. & Kondo, C. Y. Effects of septal lesions on food and water intake and operant responding for food. Physiology & Behavior 4, 729–732 (1969).

32. Stoller, W. L. Effects of septal and amygdaloid lesions on discrimination, eating and drinking. Physiology & Behavior 8, 823-IN4 (1972).

33. Bai, L. et al. Genetic Identification of Vagal Sensory Neurons That Control Feeding. Cell 179, 1129–1143.e23 (2019).

34. Madisen, L. et al. A robust and high-throughput Cre reporting and characterization system for the whole mouse brain. Nat Neurosci 13, 133–140 (2010).

35. Tong, Q., Ye, C.-P., Jones, J. E., Elmquist, J. K. & Lowell, B. B. Synaptic release of GABA by AgRP neurons is required for normal regulation of energy balance. Nat Neurosci 11, 998– 1000 (2008).

36. De Kloet, A. D. et al. A Unique “Angiotensin-Sensitive” Neuronal Population Coordinates Neuroendocrine, Cardiovascular, and Behavioral Responses to Stress. J. Neurosci. 37, 3478–3490 (2017).

37. Van Den Pol, A. N., et al. Neuromedin B and Gastrin-Releasing Peptide Excite Arcuate Nucleus Neuropeptide Y Neurons in a Novel Transgenic Mouse Expressing Strong *Renilla* Green Fluorescent Protein in NPY Neurons. J. Neurosci. 29, 4622–4639 (2009).

38. Allen Institute for Brain Science. Developing Mouse Brain Atlas. (2004).

